# High-throughput structure determination of an intrinsically disordered protein using cell-free protein crystallization

**DOI:** 10.1101/2023.12.18.571210

**Authors:** Mariko Kojima, Satoshi Abe, Tadaomi Furuta, Kunio Hirata, Xinchen Yao, Ayako Kobayashi, Ririko Kobayashi, Takafumi Ueno

## Abstract

Intrinsically disordered proteins (IDPs) play a crucial role in various biological phenomena, dynamically changing their conformations in response to external environmental cues. To gain a deeper understanding of these proteins, it is essential to identify the determinants that fix their structures at the atomic level. Here, we developed a pipeline for rapid crystal structure analysis of IDP using a cell-free protein crystallization (CFPC) method. Through this approach, we successfully demonstrated the determination of the structure of an IDP to uncover the key determinants that stabilize its conformation. Specifically, we focused on the 11-residue fragment of c-Myc, which forms an α-helix through dimerization with a binding partner protein. This fragment was strategically fused with an in-cell crystallizing protein and was expressed in a cell-free system. The resulting crystal structures of the c-Myc fragment were successfully determined at a resolution of 1.92 Å and we confirmed that they are identical to the structures of the complex with the native binding partner protein. This indicates that the environment of the scaffold crystal can fix the structure of c-Myc. Significantly, these crystals were obtained directly from a small reaction mixture (30 μL) incubated for only 72 hours. Analysis of 8 crystal structures derived from 22 mutants revealed two hydrophobic residues as the key determinants responsible for stabilizing the α-helical structure. These findings underscore the power of our CFPC screening method as a valuable tool for determining the structures of challenging target proteins and elucidating the essential molecular interactions that govern their stability.

## Introduction

Intrinsically disordered proteins (IDPs) constitute a class of proteins which are greatly influenced by their external environment, exhibiting changes in interaction and conformation modes based on the types of binding partners.(1) Identifying the determinants that stabilize the atomic-level structure of IDPs is crucial for understanding specific aspects of their biological functions. The atomic structures of IDPs have been characterized by analyzing complexes formed by IDP fragments and their binding partners.(2-7) For example, fragments of IDPs, including c-Myc and p53, were co-crystallized or fused with their native binding partners to determine the structures of the IDP region using X-ray or NMR structure analyses.(2-8) However, conventional methods lack versatility and convenience in understanding the factors contributing to fixation of the IDP structures.(6, 8-10) The libraries of IDP structures determined without the reported binding partners will improve our understanding of the essential features of IDPs.

To determine the structures of target proteins, researchers have developed crystallization tags.(11-14) One promising method involves using porous protein crystals to immobilize target proteins.(7, 15-17) However, despite its potential, the versatility of this approach is limited. Challenges persist in the design of robust scaffold crystals that maintain high diffraction quality against protein modifications and in the crystallization procedures.(7, 15) In response to these limitations, in-cell protein crystals have emerged as promising scaffolds.(18) They ensure high diffraction quality while overcoming the time-consuming aspects of crystallization screening.(11, 19-24) Notably, some proteins have been observed to undergo spontaneous crystallization within living cells.(25, 26) Advancements in X-ray diffraction data collection and analysis and the efficiency provided by automatic data processing now enables the determination of structures for various in-cell protein microcrystals.(27-36) The implementation of a rapid diffraction measurement system specialized for in-cell microcrystals is expected to significantly accelerate structure determination of target proteins. This acceleration is particularly anticipated with respect to engineering of scaffold crystals.(22-24, 37) This integration of techniques holds great promise for advancing our understanding of protein structures and associated functions.

Polyhedra crystals, which are known for their high diffraction quality, are produced following infection by the cytoplasmic polyhedral virus as shown in Figure 1a.(27) We have successfully determined the structure of a ten-amino acid miniprotein, CLN025, fused to the polyhedrin monomer (PhM), resulting in the formation of polyhedra crystals (PhC) in insect cells.(11) These findings highlight the potential of PhC as a scaffold crystal for various proteins. Additionally, we used a cell-free protein expression system to observe that PhC crystallizes rapidly.(38) By employing a cell-free expression system from wheat germ to express PhM, we determined the structure of PhC at a high resolution of 1.80 Å from sub-micron crystals obtained at a reaction scale of 100 μL within 24 hours. This is in contrast to the conventional *in vitro* crystallization system, where a scale of 1 L or more of cell culture and several months of crystallization are required.(38) These results suggest the efficiency of the cell-free protein crystallization (CFPC) method using PhC as a scaffold crystal for rapid screening systems for structural analysis.

**Fig. 1.**
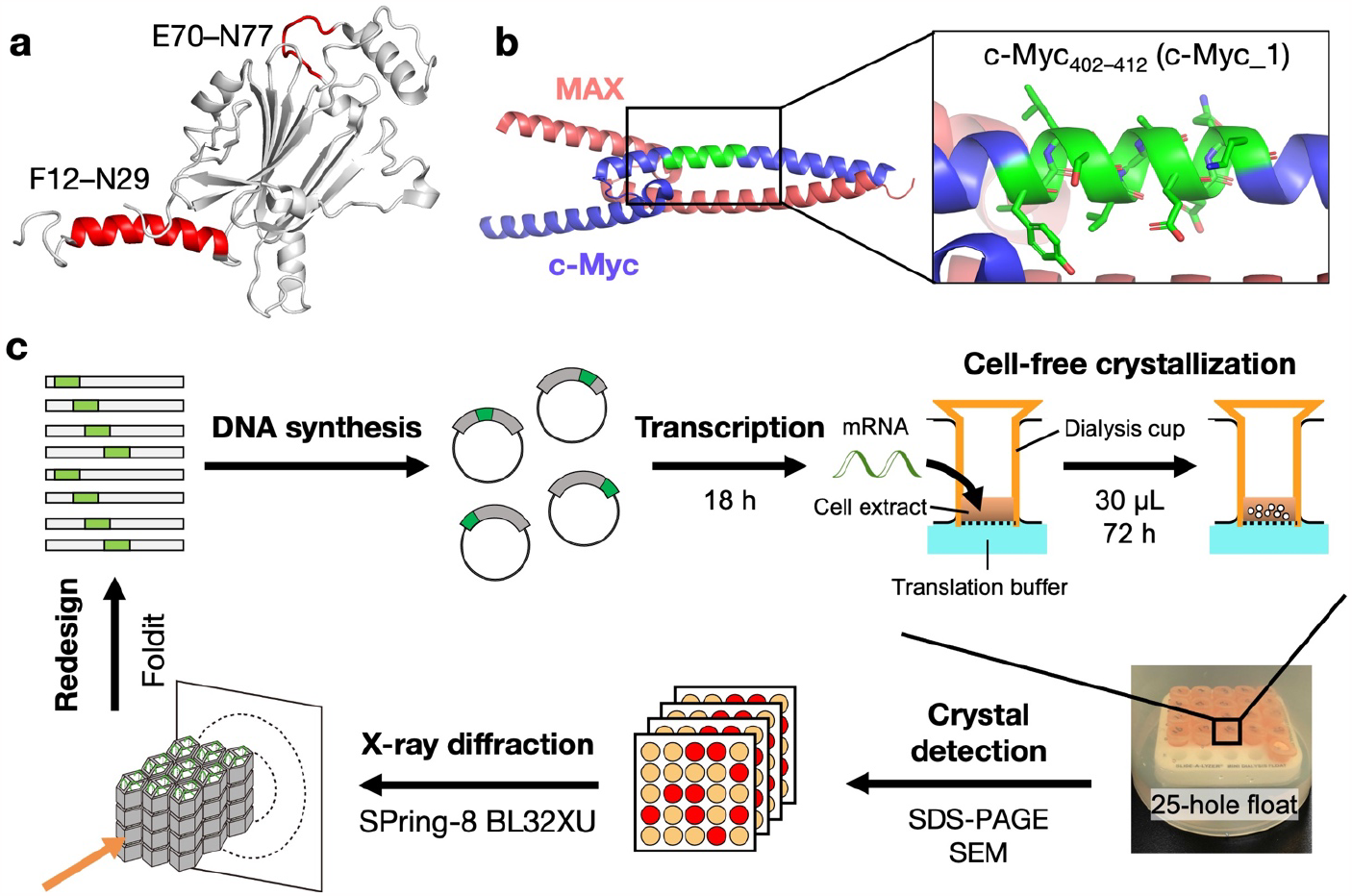
(a) Crystal structure of PhM (PDB ID: 7XWS) and two sites in PhM for fusion with c-Myc fragment_.(27)_ (b) Crystal structure of c-Myc/MAX (PDB ID: 1NKP).(8) Black square shows the position of c-Myc_402–412_ (c-Myc_1) colored green. (c) Scheme of high-throughput screening of X-ray crystallography for proteins.

In this study, we established a pipeline for rapid structure determination by integrating CFPC and Foldit, a structure prediction tool (Fig. 1).(8, 39, 40) Our goal was to elucidate the structural regulatory mechanism of the intrinsically disordered protein (IDP) region of c-Myc, a transcription factor known to bind to MAX. Previous studies have shown that the interaction between c-Myc and MAX is hindered by drug molecules binding to the short IDP region consisting of 10-11 residues (Fig. 1b).(41-47) To achieve this, we designed a candidate fusion protein using PhC as a crystal template. Using the protein structure prediction software Foldit, we identified six structures predicted to be stably crystallized (Fig. 1c). Subsequently, these structures were rapidly synthesized on a small scale (30 μL) through the CFPC method. The crystals obtained were then subjected to high-resolution structure determination of the IDP region using diffraction data acquisition at the SPring-8 synchrotron facility, using beamline BL32XU. The obtained structures were used in further redesigns to identify the key determinants fixing the structure of c-Myc structure by fragment replacement or stepwise mutation. Our method allowed us to determine 8 crystal structures out of 22 mutants, revealing that interactions of residues I403 and V406 in c-Myc contribute significantly to stabilizing the α-helical structure. This comprehensive approach enables successful determination of the IDP structure and also provides insights into the specific residues involved in stabilizing the protein’s secondary structure.

## Results

### Computational design of c-Myc fragment-fused PhMs

The present study involves rational design of the IDP fragment-fused PhM. The target fragment for structure determination was Y402–K412 of c-Myc (c-Myc_1: YILSVQAEEQK), the target sequence of pharmaceutical molecule 10058-F4.(41) It is known that c-Myc_1 is a region that undergoes conformational changes depending on the external environment, becoming an α-helix upon binding to MAX and a loop upon binding to 10058-F4. The objective was to elucidate the determinants that fix this region into the α-helix during an interaction with a neighbor protein. We selected the two regions of PhM, F12–N29 and E70–N77, as fusion sites (Fig 1a). F12–N29 is the N-terminal α-helix region in PhM which has a conformation highly similar to that of c-Myc_1 in the c-Myc/MAX complex reported previously (Fig. 1a and 1b). E70–N77 in PhM was selected as another fusion site because the ten-residue miniprotein was fixed in this region previously (Fig. 1a and 1b).(11) We designed c-Myc_1 fused PhM and optimized the fused position using Foldit. Six mutants were designed in this way and then the CFPC method was used for simultaneous crystallization of all the mutants and X-ray diffraction measurements.

We optimized the fusion position to fix the conformation of target IDP fragments in PhC by Foldit, which simulates bond formation among amino acids to calculate the increase and decrease of Rosetta energy E accompanied by the bond formation among amino acids (Table S1).(39) The tetramer of WT-PhM was used as the initial structure to design a mutant of PhM in which c-Myc_1 is fused at the F12–N29 region, which is named helix 1 (H1) (Fig. S1a). The mutant PhM in which R15–Y25 was replaced with c-Myc_1 showed the lowest E value (931.4 kcal/mol). The mutant PhMs in which E16– N26 and G14–Q24 were replaced with c-Myc_1 showed the second and the third lowest E values of 1033.0 kcal/mol and 1522.1 kcal/mol, respectively. Based on these results, mutant PhMs in which G14–Q24, R15–Y25, and E16–N26 were replaced with c-Myc_1 (**c-Myc_1/PhM**_**Δ14–24**_, **c-Myc_1/PhM**_**Δ15–25**_, and **c-Myc_1/PhM**_**Δ16–26**_, respectively) were designed (Table 1). To design a mutant PhM in which c-Myc_1 fused at the loop region of WT-PhM (E70–N77), which is named loop 1 (L1), the trimer of WT-PhM was used as the initial structure (Fig. S1b). Finally, three mutants in which Y71–D81, R72–E82, and E73–Y83 of PhM were replaced with c-Myc_1, **c-Myc_1/PhM**_**Δ71–81**_ (585.4 kcal/mol), **c-Myc_1/PhM**_**Δ72–82**_ (605.4 kcal/mol), and **c-Myc_1/PhM**_**Δ73–83**_ (627.6 kcal/mol), were designed due to the low E values (Table 1 and S1).

**Table 1.**
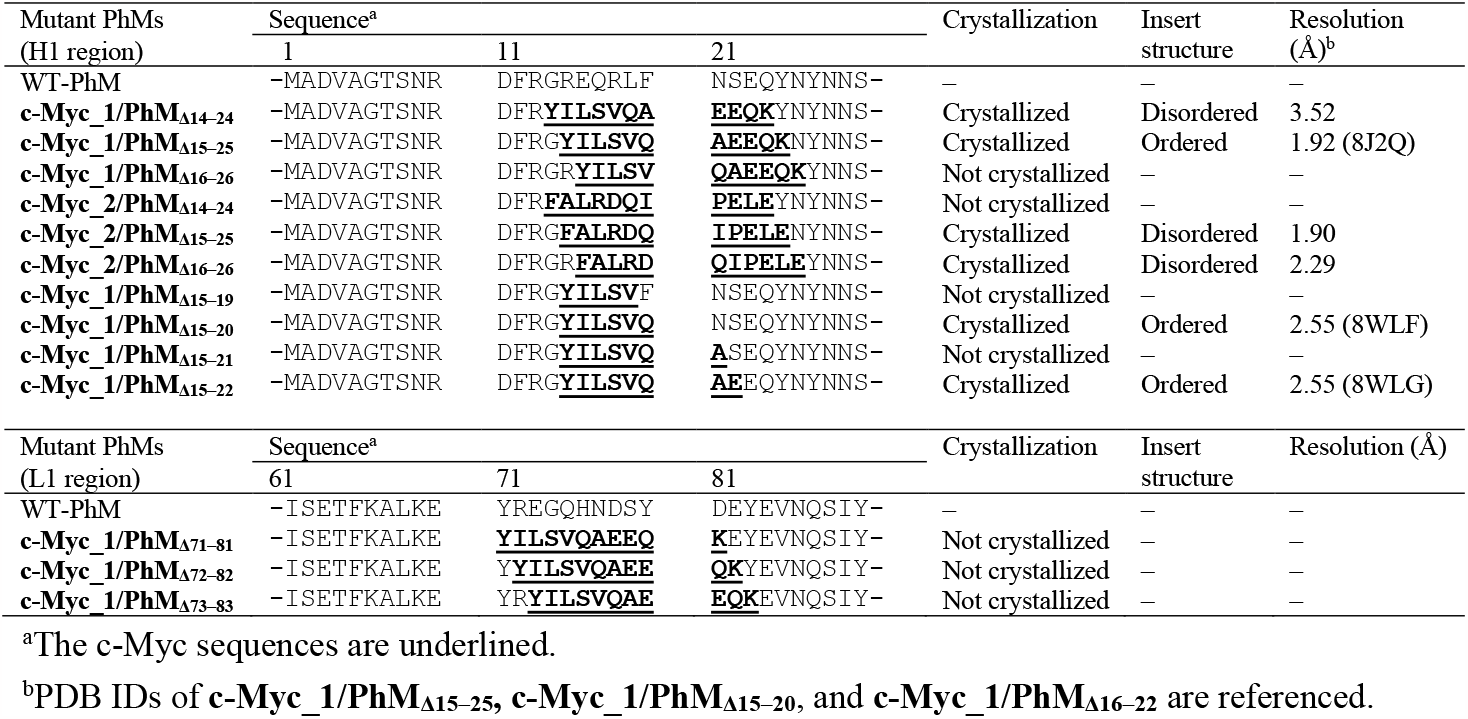
Amino acid sequences of H1 and L1 regions of WT-PhM and the designed mutant crystals of c-Myc fragment-fused PhMs.

### Crystallization and structure determination of c-Myc_1 fused PhCs

A group of six mutant PhCs were synthesized using the Wheat Germ Protein Synthesis kit (WEPRO®7240 Expression Kit). In this study, the crystallization reactions were performed in a 30 μL scale reaction cuvette fixed to a 25-hole float to carry out the crystallization of many mutants simultaneously with the minimum reaction volume that would yield crystals showing diffraction peaks sufficient for structural analysis (Fig. 1c).(38) The expression of all mutant PhMs was confirmed by SDS-PAGE (Fig. S2). The formation of the cubic crystals was observed for **c-Myc_1/PhC**_**Δ14–24**_, and **c-Myc_1/PhC**_**Δ15–25**_ using SEM (Fig. S3). The crystal structures of the mutant PhCs were determined using a micro-X-ray beam of the synchrotron SPring-8 BL32XU beamline, ZOO, and KAMO, an automated data collection and processing system.(31, 48, 49) The measurements were performed using small wedge serial crystallography (SWSX) implemented in ZOO.

The crystal structures of **c-Myc_1/PhC**_**Δ14–24**_ and **c-Myc_1/PhC**_**Δ15–25**_ were determined at resolutions of 3.52 Å and 1.92 Å, respectively (Table S2). These structures have the same *I*23 space group as WT-PhC. The unit-cell parameters (*a* = *b* = *c*) of **c-Myc_1/PhC**_**Δ14–24**_ and **c-Myc_1/PhC**_**Δ15– 25**_ are 107.91 Å and 105.97 Å, respectively. These parameters are increased relative to the cell parameter of WT-PhC (103.60 Å). The root-mean-square deviations of the Cα atoms (Cα-RMSD) of all of the mutant PhCs from WT-PhC were found to be less than 0.56 Å. The electron density corresponding to the c-Myc_1 fragment in **c-Myc_1/PhM**_**Δ15–25**_ was successfully observed between G14 and N26 (Fig. 2, S7b, and S8c). The full-length c-Myc_1 was determined based on the 2|F_o_|−|F_c_| and |F_o_|−|F_c_| electron density maps (Fig. 2b). The fragment of c-Myc_1 in **c-Myc_1/PhC**_**Δ15–25**_ (**c-Myc_1**^**Δ15-25**^) forms a helical structure which is structurally similar to the helical structure of the c-Myc/MAX complex (c-Myc^MAX^) (Fig. 2c) and the structure predicted by Foldit. The all-atom RMSD value of **c-Myc_1**^**Δ15–25**^ determined with respect to the c-Myc^MAX^ complex is 0.47 Å. This suggests that c-Myc_1 forms an α-helical structure regardless of the absence of the original interaction with the specific binding partner. The c-Myc_1 fragment in **c-Myc_1/PhC**_**Δ14–24**_ (**c-Myc_1**^**Δ14–24**^) could not be modeled because electron densities corresponding to these residues were missing (Fig. S7a and S8b).(50)

**Fig. 2.**
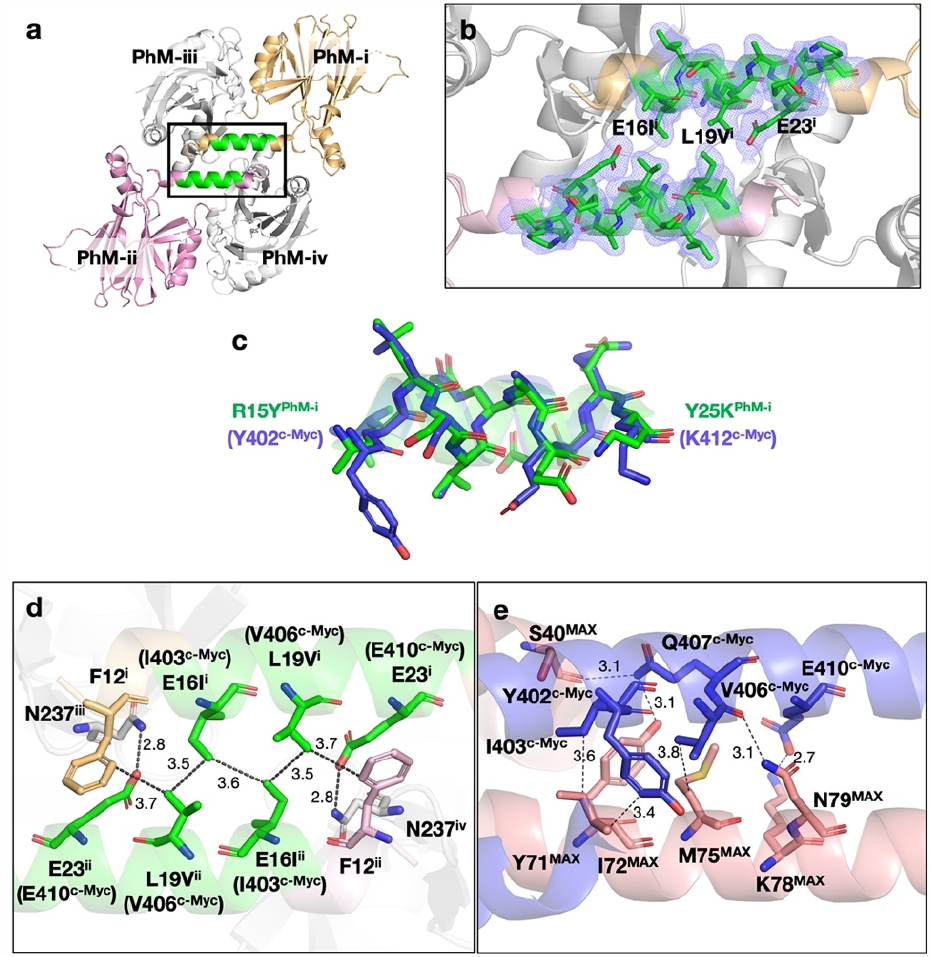
Crystal structure of **c-Myc_1/PhC**_**Δ15–25**_. (a) Structure of the tetramer of **c-Myc_1/PhC**_**Δ15–25**_. Each monomer in tetramer is named as PhM-i, PhM-ii, PhM-iii, and PhM-iv, respectively. (b) Close-up view of the c-Myc_1 fusion site of **c-Myc_1/PhC**_**Δ15–25**_. (c) Superposed structure of **c-Myc_1**^**Δ15-25**^ (green) and c-Myc^MAX^ (blue) (PDB ID: 1NKP). Noncovalent interactions between (d) **c-Myc_1**^**Δ15-25**^ and the surrounding residues in **c-Myc_1/PhC**_**Δ15–25**_, and between (e) c-Myc^MAX^ and the surrounding residues in c-Myc/MAX complex. All fragments of **c-Myc_1**^**Δ15-25**^ and c-Myc^MAX^ are colored green. The selected 2|Fo|−|Fc| electron density maps at 1.0σare shown in blue. N, O, and S atoms are colored blue, red, and yellow, respectively. The cut-off distances of noncovalent interactions are 3.5 Å and 5.75 Å for hydrogen bonds and hydrophobic interactions, respectively.(51, 52) The hydrophobic interactions of the shortest distance between two residues were shown in (d) and (e).

### Crystal structure of c-Myc_1/PhC_Δ15–25_

The **c-Myc_1**^**Δ15–25**^ fragment is located at the two-fold interfaces of the H1 helixes of two PhMs (PhM-i and PhM-ii) (Fig. 2). These helixes are known as the key domains for the crystallization of PhMs.(27) Two fragments of **c-Myc_1**^**Δ15–25**^ are directed oppositely in the middle of H1 regions. The residues E16I^i^ (I403^c-Myc^), L19V^i^ (V406^c-Myc^), and E23^i^ (E410^c-Myc^) are placed at the interface of two fragments of **c-Myc_1**^**Δ15–25**^ in an arrangement which is identical to the helix-helix conformation in c-Myc/MAX complex (Fig. 2d and 2e). **c-Myc_1**^**Δ15–25**^ of PhM-i interacts with PhM-ii via hydrophobic interactions of δ1/E16I^i^(I403^c-Myc^)–Cδ1/E16I^ii^(I403^c-Myc^) (3.6 Å), Cδ1/E16I^i^(I403^c-Myc^)–Cγ1/L19V^ii^(V406^c-Myc^) (3.5 Å) and Cγ1/L19V^i^(V406^c-Myc^)–Cδ1/F12^ii^ (3.7 Å), and with PhM-iv via the hydrogen bond of Oε2/E23^i^(E410^c-Myc^)–Nδ2/N237^iv^ (2.8 Å). The same interactions in PhM-ii also work symmetrically, since it is located in the two-fold interface (Fig. 2d). Among these interactions, I403^c-Myc^ and V406^c-^ Myc are key residues in the interactions with MAX and 10058-F4 (Fig. 2e).(8, 41) This result indicates that both the α-helical and loop conformations of **c-Myc_1**^**Δ15–25**^ are fixed by the hydrophobic interactions with I403^c-Myc^, V406^c-Myc^. This is the reason why the region of **c-Myc_1**^**Δ14–24**^ could not be determined. The fixation of α-helices in **c-Myc_1**^**Δ14–24**^ would be prevented because the hydrophobic interaction pair of I403^c-Myc^ and V406^c-Myc^ was not retained.

The sequence difference in the H1 region between WT-PhM and **c-Myc_1/PhC**_**Δ15-25**_ resulted in changes in the curvature of the helical structure. The local bending analysis of the H1 regions of them indicates that the helical region R18S^i^(S405^c-Myc^)–E23^i^(E410^c-Myc^) of the **c-Myc_1/PhC**_**Δ15-25**_ remains approximately linear, while the same region of WT-PhC has larger bending angles (Fig. S9).(53) This is caused by the change in the intermolecular interactions at the two-fold interface of H1s. The curved structure of H1 in WT-PhM is fixed by the two layers of hydrophobic and hydrophilic interaction networks (Fig. S10). The hydrophobic interaction of R15^i^–L19^ii^ observed in WT-PhM is swapped to E16I^i^(I403^c-Myc^)–L19V^ii^(V406^c-Myc^) in **c-Myc_1/PhC**_**Δ15–25**_, which induces the formation of the linear structure of H1s (Fig. 2d and S10). The change of the structures of H1s at the two-fold interface between WT-PhC and **c-Myc_1/PhC**_**Δ15–25**_ induces rearrangement of the molecular packing in the crystals. The distance of Cα/L31^i^–Cα/L31^ii^, which is located at the end of H1s, extends from 43.5 Å in WT-PhC to 46.3 Å in **c-Myc_1/PhC**_**Δ15–25**_ (Fig. S11a and S11b). This elongation indicates sliding of PhM-i and PhM-ii in opposite directions, which causes the unit-cell parameter from 103.6 Å for WT-PhC to 105.98 Å for **c-Myc_1/PhC**_**Δ15–25**_. Furthermore, some of the intermolecular interactions among M1–R10, E70–S102, Y165–V179, and N185–N196 located at the interfaces composed of three PhM trimers (PhT-A, PhT-B, and PhT-C) are broken by the sliding of PhM-i and PhM-ii (Fig. S11c). This causes structural changes of Y165–V179 and N185–N196 and induces disorder at M1–S8 and E70–S102. In addition, a structural change at Y165–V179 induces the disorder observed at A129–D134.

### Replacement of c-Myc_1 with another drug binding site of c-Myc: c-Myc_2

To investigate the specificity of the fusion site of PhC for c-Myc_1, c-Myc fragment fused PhCs in which c-Myc_1 fragments in **c-Myc_1/PhM**_**Δ14–24**_, **c-Myc_1/PhM**_**Δ15–25**_, and **c-Myc_1/PhM**_**Δ16–26**_ were replaced by another drug binding site, F375–E385 of c-Myc (c-Myc_2: FALRDQIPELE), and crystallized. The obtained cubic crystals are designated **c-Myc_2/PhC**_**Δ14-24**_, **c-Myc_2/PhC**_**Δ15–25**_, **c-Myc_2/PhC**_**Δ16–26**_, respectively (Table 1 and Fig. S4). The structures of **c-Myc_2/PhC**_**Δ15–25**_ and **c-Myc_2/PhC**_**Δ16–26**_ were determined at resolutions of 1.90 Å and 2.29 Å, respectively (Table S2). The structure of **c-Myc_2/PhC**_**Δ14-24**_ could not be determined due to insufficient diffractions. The cell parameters (a = b = c) of **c-Myc_2/PhC**_**Δ15–25**_ and **c-Myc_2/PhC**_**Δ16–26**_ were found to be106.40 Å and 106.75 Å, respectively, which are larger than that of WT-PhC (103.6 Å). The structure of the c-Myc_2 fragment in **c-Myc_2/PhC**_**Δ15–25**_ and **c-Myc_2/PhC**_**Δ16–26**_ could not be modeled because electron densities corresponding to these residues are missing (Fig. S7c, S7d, S8d, and S8e).

### MD simulations of c-Myc fragment-fused PhC

All-atom MD simulations using AMBER were performed to investigate the structures of c-Myc_1 and c-Myc_2 in PhM. The monomers **c-Myc_1/PhM**_**Δ15–25**_ and **c-Myc_2/PhM**_**Δ15–25**_ were first subjected to 100 ns MD simulations (Fig. S12). The initial structures were modeled based on the WT-PhM by replacing the original sequence with those of c-Myc_1 and c-Myc_2 fragments using PyMOL. The Cα-RMSD value of c-Myc_1 from the initial structure was retained at less than 2 Å and the structures maintain their initial helical conformations (Fig. S12a and S12c). This suggests that the α-helical structure of c-Myc_1 formed in the crystal is maintained without the interactions with binding partners in the MD simulation. The Cα-RMSD value of c-Myc_2 from the initial structure was found to vary in the range of 1-4 Å and the structure was changed to form a partially coiled structure (Fig. S12b and S12d). The difference in the converged structures of c-Myc_1 and c-Myc_2 shows that the secondary structure in monomer states is defined by the sequence of c-Myc fragments.

The effect of the molecular packing in fixing the helical structure of c-Myc_1 in the crystal environment was evaluated by performing MD simulations of tetrameric **c-Myc_1/PhC**_**Δ15–25**_ and **c-Myc_2/PhC**_**Δ15–25**_. In consideration of the crystal environment, positional restraints on backbone atoms were applied except for the c-Myc fused region. As a result, the Cα-RMSD value of c-Myc_1 from the initial structure was retained at less than 2 Å and the structure was retained as an α-helix which is the same as that of the monomer system, whereas the structure of c-Myc_2 was found to become rapidly unfolded (Fig. 3). The results of MD simulations suggest that c-Myc_1 is fixed cooperatively as a result of the structural stability of the α-helical conformation and the sufficient space around the fragment.

**Fig. 3.**
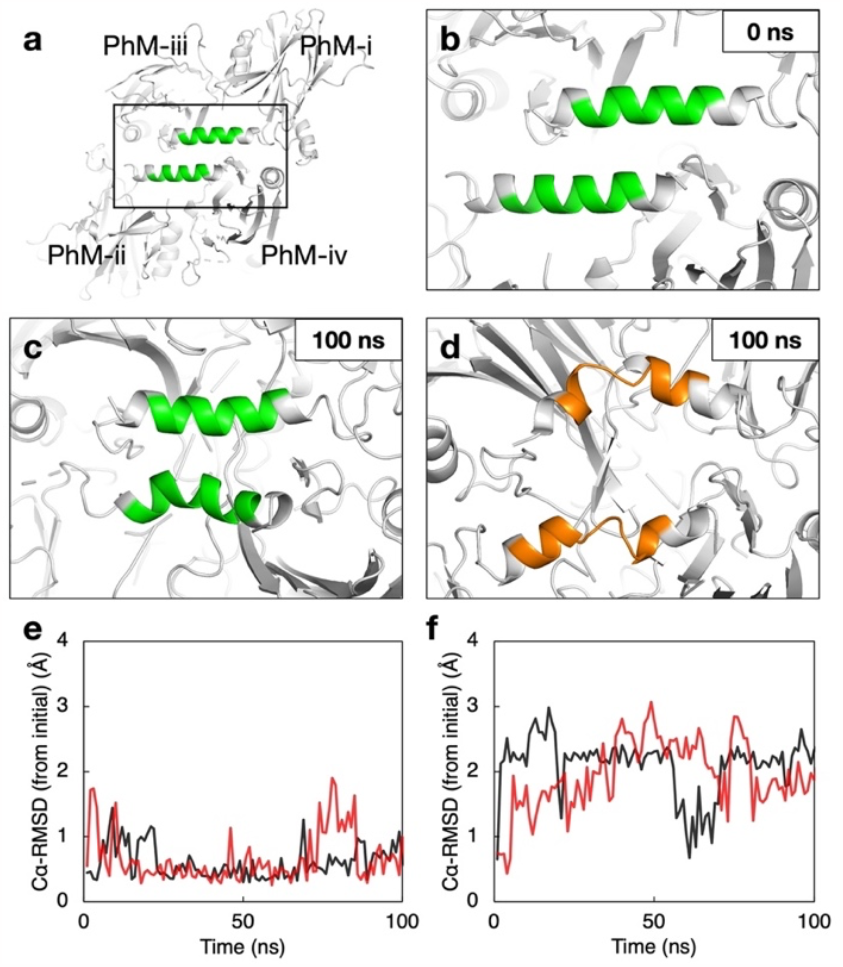
Analysis of conformations of c-Myc fragments in MD simulations of a tetramer of PhC. (a) The initial structure of tetrameric **c-Myc_1/PhC**_**Δ15–25**_. (b) The close-up view of two fragments of c-Myc_1 in the initial structure. The structures at 100 ns in MD simulations of two c-Myc fragments in (c) **c-Myc_1/PhC**_**Δ15–25**_ and (d) **c-Myc_2/PhC**_**Δ15–25**_. Time courses of the Cα-RMSD values of (e) c-Myc_1 and (f) c-Myc_2 from the initial structures. The ribbon models for c-Myc_1 in (a-c) and c-Myc_2 in (d) are colored in green and orange, respectively. Data from each monomer, PhM-i and -ii, and tetramer are colored in black and red, respectively.

### Stepwise replacement of the c-Myc_1 sequence

Given the fact that E16I^i^(I403^c-Myc^), and L19V^i^(V406^c-Myc^) are the key residues involved in formation of the hydrophobic interactions between the two α-helixes of **c-Myc_1**^**Δ15–25**^, we designed the mutants of **c-Myc_1/PhC**_**Δ15–25**_, in which F20Q–Y25K, N21A–Y25K, S22E–Y25K, and Y25K were replaced with the original sequence of WT-PhM. These mutants are designated **c-Myc_1/PhM**_**Δ15–19**_, **c-Myc_1/PhM**_**Δ15–20**_, **c-Myc_1/PhM**_**Δ15–21**_, and **c-Myc_1/PhM**_**Δ15–22**_, respectively (Table 1). Four PhM mutants were synthesized in the above-mentioned process (Fig. S2d). **c-Myc_1/PhM**_**Δ15–20**_ and **c-Myc_1/PhM**_**Δ15–22**_ were successfully crystallized and designated **c-Myc_1/PhC**_**Δ15–20**_ and **c-Myc_1/PhC**_**Δ15–22**_ as crystals, respectively, whereas others were not successfully crystallized (Fig. S5). The crystal structures of both crystals were determined at a resolution of 2.55 Å (Tables 1 and S2).

Despite the replacement of the sequence, the structures of the c-Myc_1 fragment in **c-Myc_1/PhC**_**Δ15–20**_ and **c-Myc_1/PhC**_**Δ15–22**_ were found to be identical to that of **c-Myc_1**^**Δ15–25**^ (Fig. 4, S7e, S7f, S8f, and S8g). Hydrophobic interactions at E16I^i^(I403^c-Myc^) and L19V^i^(V406^c-Myc^) were also formed both in **c-Myc_1/PhC**_**Δ15–20**_ and **c-Myc_1/PhC**_**Δ15–22**_, which enhances that these two hydrophobic residues are essential to fix the α-helical structure of c-Myc_1 fragment (Fig. 4a, 4c, and 4e). The hydrogen bond at Oε2/E23^i^(E410^c-Myc^)–Nδ2/N237^iv^ was not observed in **c-Myc_1/PhC**_**Δ15–22**_. This suggests that this hydrogen bond is not essential but supports the stability of the structure of c-Myc fragment (Fig. 4e). We evaluated the stability of the backbone structure of the c-Myc_1 fragment in each mutant PhC based on the normalized mean B-factor (B’) (Fig. 4g and S13). The B’ value of R15Y(Y402^c-Myc^)–Y25 in **c-Myc_1/PhC**_**Δ15–22**_ is higher than in **c-Myc_1**^**Δ15–25**^. This indicates that the c-Myc_1 fragment in **c-Myc_1/PhC**_**Δ15–22**_ vibrates due to a lack of the hydrogen bonds at E23^i^ (Fig. 4e and 4g).

**Fig. 4.**
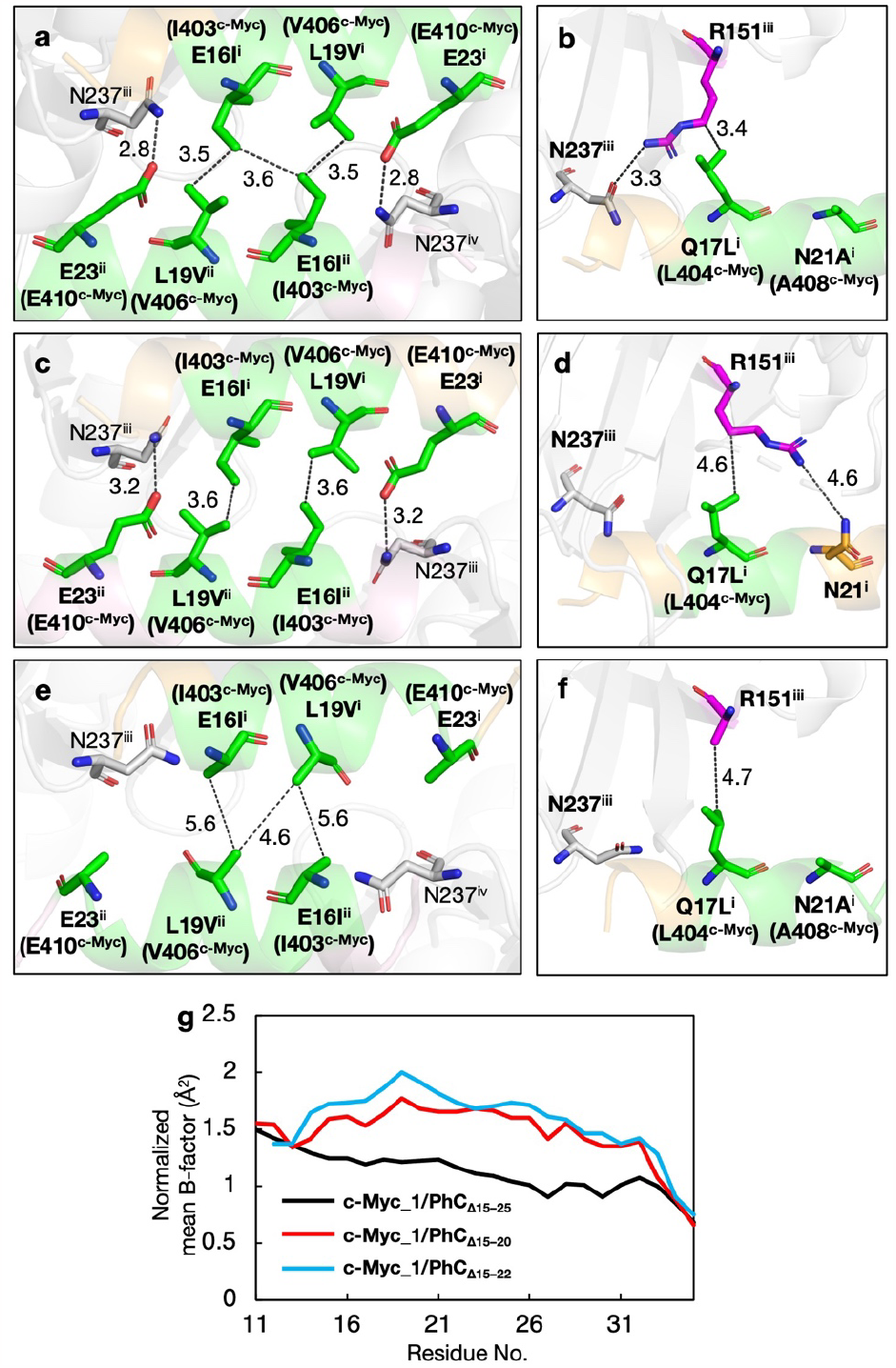
Structure analysis of **c-Myc_1/PhM**_**Δ15–25**_, **c-Myc_1/PhM**_**Δ15–20**_, and **c-Myc_1/PhM**_**Δ15–22**_. The intermolecular interaction networks of two-fold interface of c-Myc_1 fragments of (a) **c-Myc_1/PhM**_**Δ15–25**_, (c) **c-Myc_1/PhM**_**Δ15–20**_, and (e) **c-Myc_1/PhM**_**Δ15–22**_. Intermolecular interactions at R151^iii^ of (b) **c-Myc_1/PhM**_**Δ15–25**_, (d) **c-Myc_1/PhM**_**Δ15–20**_, and (f) **c-Myc_1/PhM**_**Δ15–22**_. (g) The spectrum of the normalized B factor (B’) corresponding to D11–V35 of **c-Myc_1/PhM**_**Δ15–25**_, **c-Myc_1/PhM**_**Δ15–20**_, and **c-Myc_1/PhM**_**Δ15–22**_. E16I^i^(I403^c-Myc^), L19V^i^(V406^c-Myc^), and E23^i^(E410^c-Myc^) in (e) are displayed as alanine due to lack of the corresponding electron densities of the sidechain. All fragments of c-Myc_1 are colored green. R151^iii^ in (b), (d), and (f) are colored magenta. The cut-off distances of noncovalent interactions are 3.5 Å for hydrogen bonds, and 5.75 Å for hydrophobic interactions.(51, 52)

The increase of the B’ values of R15Y(Y402^c-Myc^)–Y25 in **c-Myc_1/PhC**_**Δ15–20**_ is caused by a lack of hydrophobic interactions at Cδ1/E16I^i^(I403^c-Myc^)–Cδ1/E16I^ii^(I403^c-Myc^) and a weakening of a hydrogen bond with PhM^iii^ (Fig. 4c and 4g). Remarkably, one of the residues surrounding the c-Myc_1 fragment, R151^iii^, undergoes a conformational change relative to **c-Myc_1/PhC**_**Δ15–25**_ (Fig. 4b and 4d). R151^iii^ in **c-Myc_1/PhC**_**Δ15–25**_ forms a hydrogen bond of Nη2/R151^iii^–Oδ1/N237^iii^ (3.3 Å) and a hydrophobic interaction of Cδ/R151^iii^–Cδ2/Q17L^i^(L404^c-Myc^) (3.4 Å), whereas only a hydrophobic interaction at Cγ/R151^iii^–Cδ2/Q17L^i^(L404^c-Myc^) (4.6 Å) is formed in **c-Myc_1/PhC**_**Δ15–20**_ around c-Myc_1 region due to the flip of R151^iii^ caused by the hydrophilic side chain of N21^i^ (Fig. 4b and 4d). The side chain of R151^iii^ in **c-Myc_1/PhC**_**Δ15–20**_ could not be modeled due to the missing electron densities. These flips of R151^iii^ destabilize the backbone structure of c-Myc_1 fragment in **c-Myc_1/PhC**_**Δ15–20**_.

To investigate the conformation of R151 in response to the sequence of c-Myc fragment, we attempted to crystallize and analyze the structures of R151 mutants of **c-Myc_1/PhC**_**Δ15–25**_, in which R151 is replaced by other residues such as Q, K, P, V, S, Y, I, and A (Fig. S2e and S6). Two PhC mutants, **c-Myc_1/PhC**_**Δ15–25**__**R151Q** and **c-Myc_1/PhC**_**Δ15–25**__**R151K** were obtained, and structures were determined at resolutions of 2.00 Å and 2.04 Å, respectively (Fig. S7g, S7h, S8h, S8i, and S14). R151Q^iii^ forms the weak hydrophobic interaction of Cγ/R151^iii^–Cδ2/Q17L^i^(L404^c-Myc^) (3.9 Å) in **c-Myc_1/PhC**_**Δ15–25**__**R151Q** (Fig. S14b). B’ values of the c-Myc_1 fragment in **c-Myc_1/PhC**_**Δ15– 25**__**R151Q** are lower than those of **c-Myc_1/PhC**_**Δ15–20**_ and higher than those of **c-Myc_1/PhC**_**Δ15–25**_ (Fig. S13). B’ values of c-Myc_1 fragment in **c-Myc_1/PhC**_**Δ15–25**__**R151K** are lower than those of **c-Myc_1/PhM**_**Δ15–20**_ whereas the interactions between R151K^iii^ and c-Myc_1 fragment are weak. The side chain of R151K^iii^ could not be modeled due to missing electron densities (S14d). This result suggests that the rigid hydrogen bond of Oε2/E23^i^(E410^c-Myc^)–Nδ2/N237^iv^ (2.9 Å) overcomes the destabilization of the α-helical structure of the c-Myc_1 fragment (Fig. S14c). Therefore, the stability of the c-Myc_1 fragment is reinforced by the hydrogen bond of Oε2/E23^i^(E410^c-Myc^)–Nδ2/N237^iv^ and the hydrogen bond of R151^iii^.

## Discussion

Rapid screening using CFPC was established in our process for determining IDP fragments. The structure of the 11-residue IDP fragment was successfully determined at a suitable site predicted with Foldit. In all, 22 of PhMs fused with c-Myc_1 or c-Myc_2 fragments were investigated within one month. Among these PhMs, 8 were found to diffract with sufficiently high resolution to analyze the structures with the microfocus X-ray beamline and 5 were determined as the ordered structure of the c-Myc_1 fragment. We optimized the fusion position of c-Myc_1 in PhM by Foldit to estimate the potency of the structure determination and obtained a high-resolution structure of **c-Myc_1**^**Δ15–25**^. This result suggests that the predicted structures computed by the versatile software can be quickly evaluated using our CFPC screening process. The trial numbers of crystallization are expected to be improved by the automated system for DNA construction and protein expression in the future.(54-57) An automated screening system can be applied for comprehensive analysis, such as an alanine scan, to establish the crystal design strategy. Crystal design will be diversified by using composite crystals of proteins and organic compounds as scaffold crystals.(58) The information obtained by screening is expected to be used as training data for deep learning.(59) By combining the deep learning-based design and construction of a library of tailored fusion protein crystals, the target molecules of our screening will be extended to various proteins, peptides, and small organic molecules.

In this study, the structure determination of a pharmaceutical binding site of c-Myc was conducted. As a result of the insertion of two sequences of c-Myc fragment, c-Myc_1 and c-Myc_2, to PhM, the structure of c-Myc_1 was determined as **c-Myc_1**^**Δ15-25**^ which has a structure identical to that of c-Myc^MAX^. **c-Myc_1**^**Δ15-25**^ structure is fixed by the intermolecular interaction network between two H1s which are essential interactions. This observation agrees with the finding that the c-Myc_1 fragment in **c-Myc_1/PhM**_**Δ14–24**_ could not be determined because of the mismatch of the inter-helix interactions. Stepwise replacement of the c-Myc_1 sequence with the original sequence of PhM revealed that the hydrophobic interactions at E16I^i^ (I403^c-Myc^) and L19V^i^ (V406^c-Myc^) are required to fix the helical structure of c-Myc_1 fragment in PhC. Our result is the first report which identifies the interactions required to fix the helical conformation of c-Myc in the experiments. Thus, we demonstrated that the pipeline consisting of computational design, multi-sample CFPC, and automated X-ray diffraction measurements achieves high-throughput screening for high-resolution structure analysis, consequently identifying the intermolecular interactions needed to fix the IDP structures by stepwise amino acid replacement.

Furthermore, these results serve as a model for structure determination of protein complexes. Symmetric molecular interfaces in PhC can be utilized as fusion sites for the target protein, which forms oligomers. Asymmetric interfaces in protein crystals serve as fusion sites for asymmetric protein complexes, such as protein-ligand and IDP-interaction partner complexes. Structure determination of the various structures of one IDP requires searching for and designing synthetic ligands and binding partner proteins. To fix the c-Myc in other structures, the interactions stabilizing the helical structure of c-Myc should be inhibited. We previously reported that a ten-amino acid miniprotein which folded into a β-hairpin structure in a steady state was folded into the loop-helix-loop structure in a metastable state by salt bridges between the target and PhC. This suggests that the noncovalent bonds change the folding energy landscape of targets. We expect to apply this strategy to structure determination of IDPs in different states fixed by surrounding proteins. The reported simulation indicates that the free energy penalty involved in changing the helical monomers of the peptide of IDP with ten amino acids to provide the other conformation is more than 7–8 kcal/mol.(60) This is approximately equal to twice the OH---O hydrogen bonding energy.(61) This means that two or more hydrogen bonds should be introduced between the target IDP and PhC in addition to energy equivalent to the force required to disrupt the original interactions to provide the other structure in PhC. Therefore, our CFPC screening system using PhC facilitates rational molecular design to clarify the number and position of the intermolecular interactions surrounding IDPs.

In conclusion, the rapid screening of crystallization of IDP fragments fused to PhM was demonstrated using the CFPC method. The whole structure of **c-Myc_1**^**Δ15-25**^ was determined by screenings combined with computational protein design. The obtained structure of **c-Myc_1**^**Δ15-25**^ is highly similar to the α-helical structure of c-Myc^MAX^ observed in the co-crystal with the original binding partner. The stepwise replacement of c-Myc_1 revealed the key intermolecular interactions to fix **c-Myc_1**^**Δ15-25**^ with an α-helix conformation as the hydrophobic interactions at E16I^i^(I403^c-Myc^) and L19V^i^(V406^c-Myc^). These results prove that our screening system is valuable for rapid structure determination of IDPs at high resolution and in determining the key residues required to fix their structures. Our screening system will be applied to target IDPs whose binding partners have not yet been identified and to design the new binding molecules such as inhibitors. Furthermore, the large number of crystal structures accumulated by rapid screening is expected to be used in construction of a design library of protein crystals, accelerating the elucidation of the mechanism of IDP folding supported by other proteins.

## Materials and Methods

Detailed Materials and Methods are available in SI Appendix.

## Acknowledgments

This work was supported by JSPS KAKENHI Grant No. JP19H02830, JP20K21244, and Grant-in-Aid for Scientific Research on Innovative Areas “Molecular Engines” (JP18H05421) to T. U. and JP18K05140 to S. A., and the Adaptable and Seamless Technology Transfer Program through Target-driven R&D (JPMJTR20U1) from the Japan Science and Technology Agency to T. U. Synchrotron radiation experiments were conducted under the approval of 2021A2772, 2021B2772, 2022A2771, and 2022B2771 at SPring-8. This work was supported by the SUNBOR Grant from the Suntory Foundation for Life Sciences to M. K. This research was partially supported by the Platform Project for Supporting Drug Discovery and Life Science Research (Basis for Supporting Innovative Drug Discovery and Life Science Research (BINDS)) from the AMED under Grant number JP21am0101070 (support number 1854).

