## Supporting information for "High-throughput structure determination of an intrinsically disordered protein using cell-free protein crystallization"

### Table of Contents


| | |
|---|---|
| **Experimental Procedures** | |
| Materials | 3 |
| Cell-free synthesis and crystallization of mutant PhCs | 3 |
| Scanning Electron Microscopy (SEM) analysis | 3 |
| X-ray crystal structure analysis | 3 |
| Molecular dynamics simulation | 4 |
| **Fig. S1** | 6 |
| **Fig. S2** | 7 |
| **Fig. S3** | 8 |
| **Fig. S4** | 9 |
| **Fig. S5** | 10 |
| **Fig. S6** | 11 |
| **Fig. S7** | 12 |
| **Fig. S8** | 13 |
| **Fig. S9** | 14 |
| **Fig. S10** | 15 |
| **Fig. S11** | 16 |
| **Fig. S12** | 17 |
| **Fig. S13** | 18 |
| **Fig. S14** | 19 |
| **Table S1** | 20 |
| **Table S2** | 21 |
| **References** | 22 |

### Experimental Procedures

#### Materials

The competent cells of DH5α *E. coli* were purchased from Toyobo and Invitrogen™ (Life Technologies). The cloning was implemented using In-Fusion® HD Cloning Kit (Takara Bio). The PCR reactions were implemented using KOD Plus Mutagenesis Kit (Toyobo). All oligo-DNA fragments for IDP fragments are purchased from GenScript and Thermo Fischer Scientific. Other reagents were purchased from TCI, Wako, Nacalai Tesque, Sigma–Aldrich, and Life Technologies and were used without further purification.

#### Cell-free synthesis and crystallization of mutant PhCs

Expression and crystallization of PhCs were performed using a WEPRO7240 Expression Kit (CellFree Sciences). The gene of PhM was cloned into the PEU-E01-MCS vector (CellFree Sciences) for the PhM expression. The plasmid DNAs of c-Myc fragment fused PhM were prepared by in-Fusion or inverse PCR using wild-type PhM in PEU-E01-MCS vector as a template. The plasmid was amplified by PCR. Transcription was performed for PCR products according to the protocol of the expression kit. After the incubation at 37 °C for 18 h, mRNA was used for translation. The translation reaction was carried out using the dialysis method. 32 μL reaction mixture containing 8 μL of WEPRO®7240 wheat germ extract, 8 μL of the mRNA solution, 40 mg/mL creatine kinase, and 16 μL of SUB-AMIX® SGC solution was added in dialysis cup and set on the 1.5 mM tubes containing 1.1 mM SUB-AMIX® SGC solution at 20 °C for 72 h. For large numbers of samples as 1st screening, those dialysis cups were arranged in float put on 60 mL of SUB-AMIX® SGC solution, and incubated at 20 °C for 72 h. White precipitates were collected after centrifuging the reaction solution and washed with PBS several times. For small-scale screening after 1st screening, the reaction volume in the dialysis cup can be increased up to a 160 μL reaction mixture as appropriate for various characterizations.

#### Scanning Electron Microscopy (SEM) analysis

The morphologies of purified crystals were confirmed by scanning electron microscopy (SEM). After substituting PBS with Milli-Q water, the crystals were dried and observed by SEM. SEM analyses were performed on JCM-6000 Neoscope™ (JEOL) and JSM-IT100 InTouchScope™ (JEOL).

#### X-ray crystal structure analysis

Before the data collection, the crystals were immersed in Grace medium containing 50% (v/v) ethylene glycol, spread over MicroMesh, and frozen in liquid nitrogen. The X-ray diffraction data of each crystal were collected at 100K at the beamline BL32XU at SPring-8 using an X-ray wavelength of 1.00 Å. The whole data collection process was automated by *ZOO* system including sample exchange by the robot.(1) X-ray diffraction data of **c-Myc_1/PhC$_{\Delta 14–24}$**, **c-Myc_1/PhC$_{\Delta 15–25}$**, **c-Myc_2/PhC$_{\Delta 15–25}$**, **c-Myc_2/PhC$_{\Delta 16–26}$**, **c-Myc_1/PhC$_{\Delta 15–22}$**, **c-Myc_1/PhC$_{\Delta 15–20}$**, **c-Myc_1/PhC$_{\Delta 15–25}$_R151Q**, and **c-**

**Myc_1/PhC$_{\Delta15–25}$_R151K** were collected at 100 K at beamline BL32XU at SPring-8 using X-ray wavelength of 1.00 Å. The complete sets of structure factor amplitudes were obtained by merging multiple small-wedge (5° or 10° each) datasets collected from single crystals. The crystal positions in a cryoloop were identified by low-dose raster scan. The whole data collection process was automated by ZOO system including sample exchange by a robot. Collected datasets were automatically processed and merged by KAMO.(2) Each dataset was indexed and integrated using XDS.(3) The datasets consistently indexed with the known cell parameter (a ~ 103 Å, I23) were selected and two possible reindex operators (hkl and -hlk) were tested to give better match to the previously solved data (2OH6). The datasets were subjected to hierarchical clustering by pairwise correlation coefficient of intensities. The datasets in each cluster were scaled and merged using XSCALE(3) with outlier rejections implemented in KAMO.(3) The clusters with the highest CC1/2 were chosen for downstream analyses. The structure was solved by rigid body refinement with phenix.refine using the previously solved structure (PDB ID: 2OH6).(4) Refinement of the protein structure was performed at resolutions of 3.52 Å, 1.92Å, 1.90 Å, 2.29 Å, 2.55 Å, 2.55 Å, 2.00 Å and 2.04 Å for **c-Myc_1/PhC$_{\Delta14–24}$**, **c-Myc_1/PhC$_{\Delta15–25}$**, **c-Myc_2/PhC$_{\Delta15–25}$**, **c-Myc_2/PhC$_{\Delta16–26}$**, **c-Myc_1/PhC$_{\Delta15–20}$**, **c-Myc_1/PhC$_{\Delta15–22}$**, **c-Myc_1/PhC$_{\Delta15–25}$_R151Q**, and **c-Myc_1/PhC$_{\Delta15–25}$_R151K**, respectively, using REFMAC5 in the of CCP4 suite.(4) Rebuilding was performed using COOT based on sigma-A weighted 2|$F_o$|-|$F_c$| and |$F_o$|-|$F_c$| electron density maps. M1–G33, K66–K105, N128–H141, Y172–I175, S187–N190, A243-Q248 in **c-Myc_1/PhM$_{\Delta14–24}$**, M1–R10, E70–S102, N128–D134, G192–A194and S245–Q248 in **c-Myc_1/PhM$_{\Delta15–25}$**, M1–Q24, E70–S102, A129–P133, S187–V191, and S245–Q248 in **c-Myc_2/PhC$_{\Delta15–25}$**, M1–N32, A67–A104, A129–S131, H170, P186–H195, and S245–Q248 in **c-Myc_2/PhC$_{\Delta16–26}$**, M1–R10, K69–N103, V130–D134, N190–A194, and S245–Q248 in **c-Myc_1/PhM$_{\Delta15–20}$**, M1–D11, K69–A104, N128–E135, E171–N173, P186–H195, and S245–Q248 in **c-Myc_1/PhM$_{\Delta15–22}$**, M1–R10, E70–N103, A129–D134, S193–A194 and S245–Q248 in **c-Myc_1/PhC$_{\Delta15–25}$_R151Q**, M1–D11, L68–A104, G127–E135, H170–N173, and G192–A194, and L244–Q248 in **c-Myc_2/PhC$_{\Delta15–25}$_R151K** could not be modeled because electron densities corresponding to these residues were missing.(5) The models were subjected to quality analysis during the various refinement stages with omit maps and RAMPAGE.(6) The diffraction and refinement statistics are summarized in Table S1. Atomic coordinates for **c-Myc_1/PhC$_{\Delta15–25}$**, **c-Myc_1/PhC$_{\Delta15–20}$**, **c-Myc_1/PhC$_{\Delta15–22}$**, **c-Myc_1/PhC$_{\Delta15–25}$_R151Q**, and **c-Myc_1/PhC$_{\Delta15–25}$_R151K** have been deposited in the Protein Data Bank under accession codes 8J2Q, 8WLF, 8WLG, 8X8V, and 8X8S, respectively. All images were produced using PyMOL (https://pymol.org/).

**Molecular dynamics simulation**

The initial structures of the monomer of **c-Myc_1/PhM$_{\Delta15–25}$** and **c-Myc_2/PhM$_{\Delta15–25}$** were constructed from the crystal structure of WT-PhC (PDB ID: 7XWS) by replacing the original sequence with those of c-Myc_1 and c-Myc_2 fragments using PyMOL. The initial structures of the tetramer of **c-Myc_1/PhM$_{\Delta15–25}$** and **c-Myc_2/PhM$_{\Delta15–25}$** were constructed from the crystal structure of c-

**Myc_1/PhM$_{\Delta15–25}$** (PDB ID: 8J2Q). The tetramer structure of **c-Myc_2/PhM$_{\Delta15–25}$** was constructed by replacing the original sequence of **c-Myc_1/PhM$_{\Delta15–25}$** using PyMOL. The nucleotides were removed from all structures. Counter ions ($Na^+$ and $Cl^-$) were added to preserve electric neutrality. For each system, the energy minimization of 300 steps was carried out with restraints on heavy atoms. Then, 500 ps equilibration under NVT condition (300 K) and 500 ps equilibration under NPT condition (300 K and 1 bar) were conducted with the same restraints above. Finally, 100 ns production runs were conducted for all systems. In tetramer systems, position restraints on backbone atoms with a force constant of 500 kcal $mol^{-1}$ $Å^{-2}$ were applied except for the c-Myc fused region, N26–Q248. All the MD simulations were performed using the Amber ff19SB force field and TIP3P water model.(7, 8) The temperature and pressure were regulated by the Berendsen thermostat and barostat, respectively. The time step was set to 2 fs, and the trajectories were recorded every 1 ns. The MDTraj software was used to analyze the RMSD.(9) PyMOL was used for visualization of the structures.

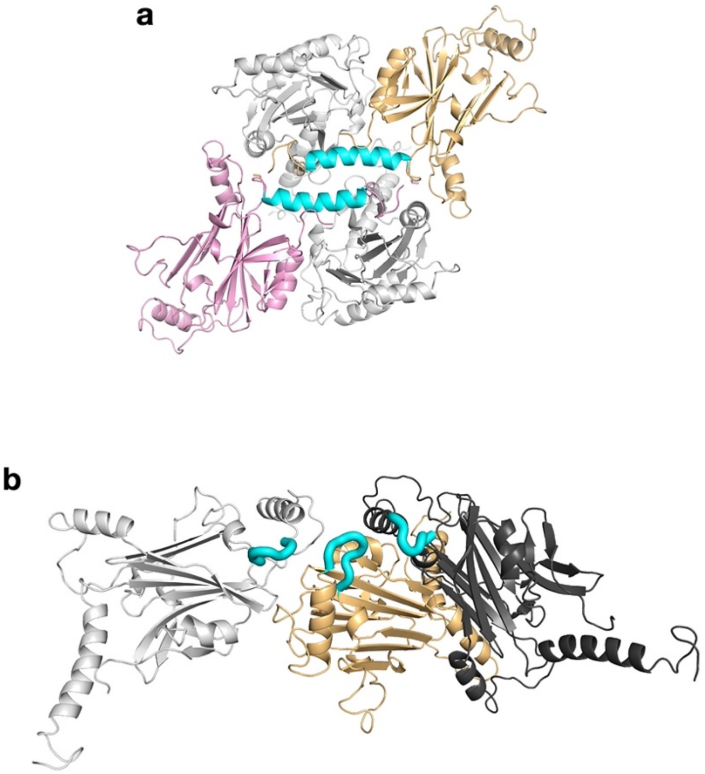


**Fig. S1** Initial structure of Foldit. (a) Tetramer and (b) trimer of wild-type PhM (WT-PhM) (PDB ID: 7XWS). The fusion sites of c-Myc fragments, H1 in (a) and L1 in (b), are colored in cyan. Proteins are represented by ribbon model.

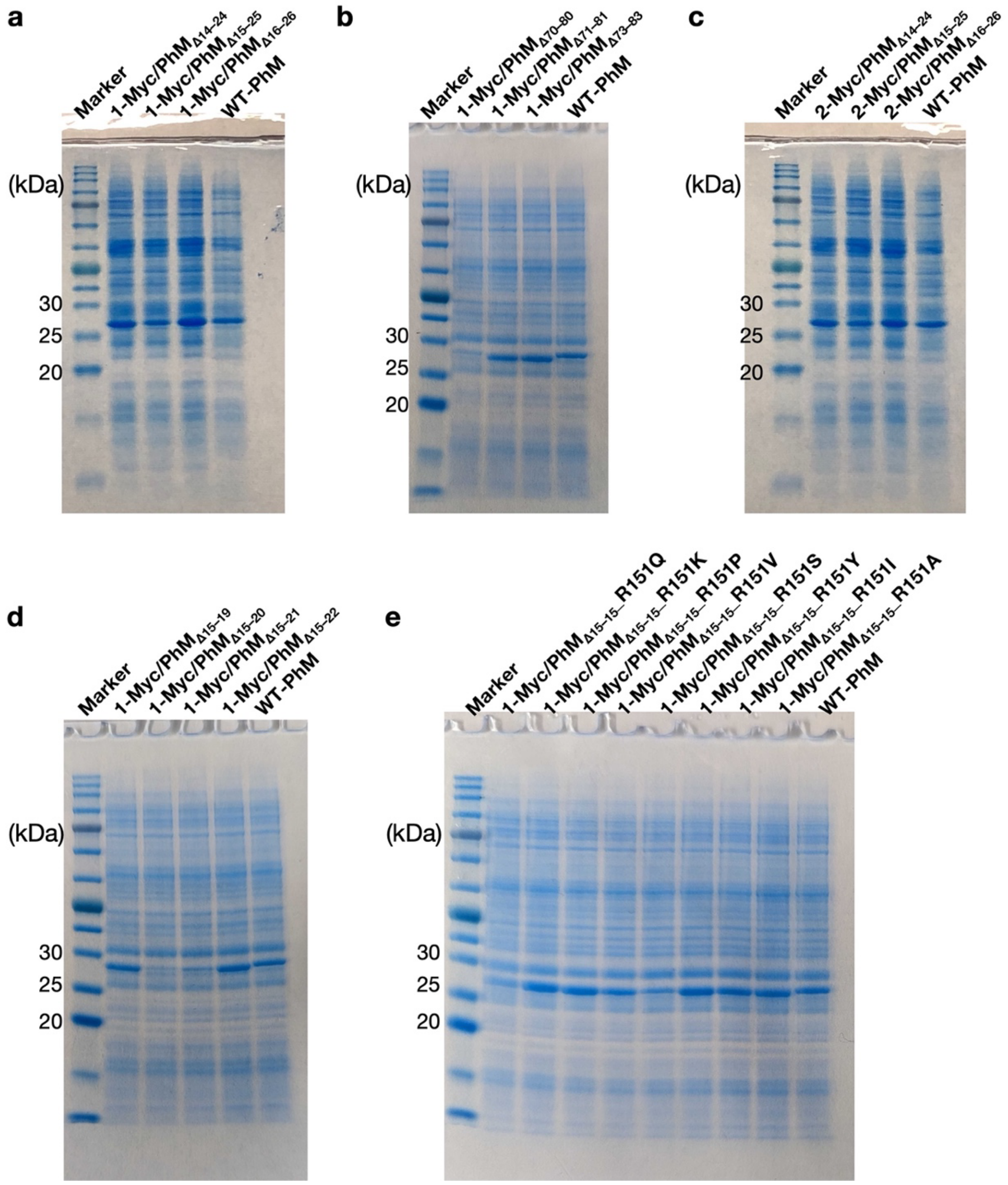


**Fig. S2** SDS-PAGE of c-Myc_1 and c-Myc_2 fragments fused PhCs; (a) **c-Myc_1/PhM$_{\Delta 14–24}$, c-Myc_1/PhM$_{\Delta 15–25}$, c-Myc_1/PhM$_{\Delta 16–26}$,** (b) **c-Myc_1/PhM$_{\Delta 70–80}$, c-Myc_1/PhM$_{\Delta 71–81}$, c-Myc_1/PhM$_{\Delta 73–83}$**, (c) **c-Myc_2/PhM$_{\Delta 14–24}$, c-Myc_2/PhM$_{\Delta 15–25}$**, and **c-Myc_2/PhM$_{\Delta 16–26}$**, (d) **c-Myc_1/PhM$_{\Delta 15–19}$, c-Myc_1/PhM$_{\Delta 15–20}$**, **c-Myc_1/PhM$_{\Delta 15–21}$**, and **c-Myc_1/PhM$_{\Delta 15–22}$**, and (e) **c-Myc_1/PhM$_{\Delta 15–25}$_R151Q, R151K, R151P, R151V, R151S, R151Y, R151I,** and **R151A.**

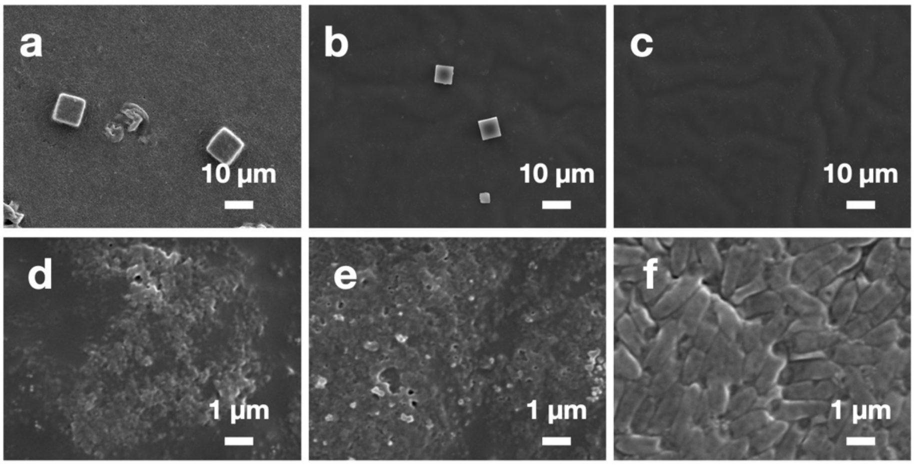


**Fig. S3** SEM images of (a) **c-Myc_1/PhC$_{\Delta14–24}$**, (b) **c-Myc_1/PhC$_{\Delta15–25}$**, (c) **c-Myc_1/PhC$_{\Delta16–26}$**, (d) **c-Myc_1/PhM$_{\Delta70–80}$**, (e) **c-Myc_1/PhM$_{\Delta71–81}$**, and (f) **c-Myc_1/PhM$_{\Delta73–83}$**.

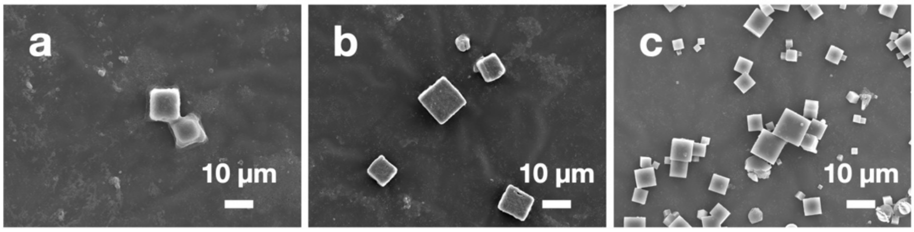

**Fig. S4** SEM images of (a) **c-Myc_2/PhC$_{\Delta14–24}$**, (b) **c-Myc_2/PhC$_{\Delta15–25}$**, and (c) **c-Myc_2/PhC$_{\Delta16–26}$**.

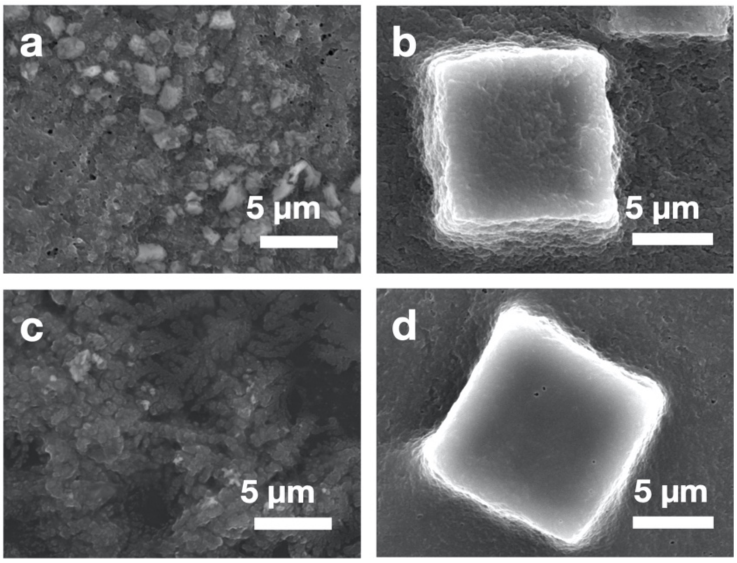


**Fig. S5** SEM images of (a) **c-Myc_1/PhM$_{\Delta 15-19}$**, (b) **c-Myc_1/PhC$_{\Delta 15-20}$**, (c) **c-Myc_1/PhM$_{\Delta 15-21}$**, and (d) **c-Myc_1/PhC$_{\Delta 15-22}$**.

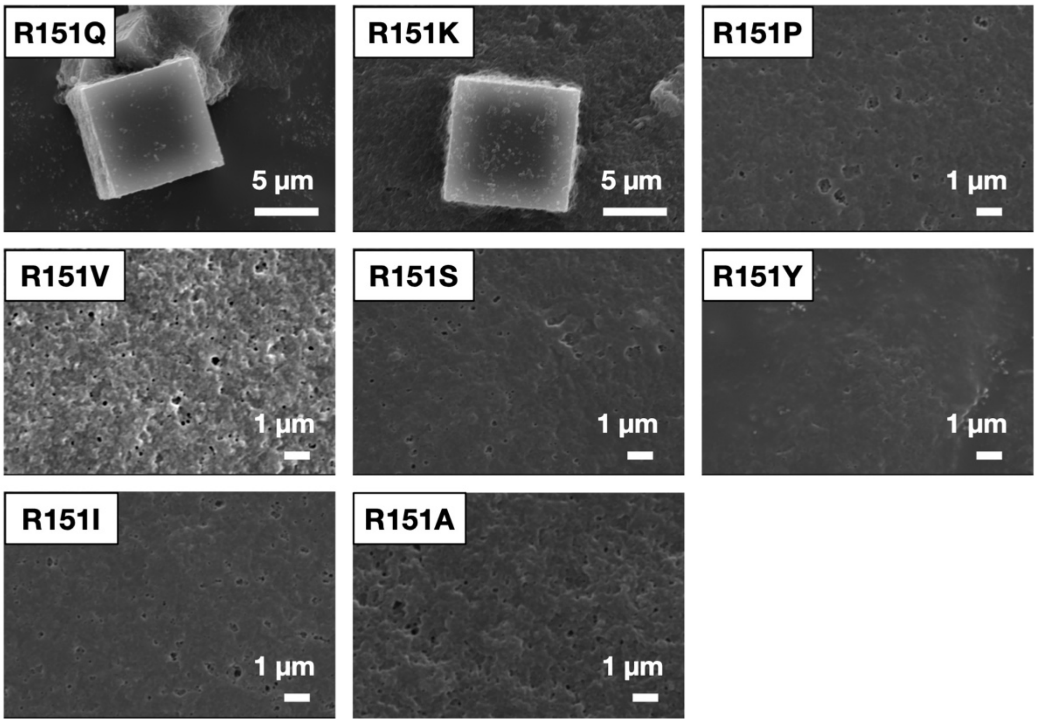

**Fig. S6** SEM images of R151 mutants of **c-Myc_1/PhC$_{\Delta 15-25}$**.

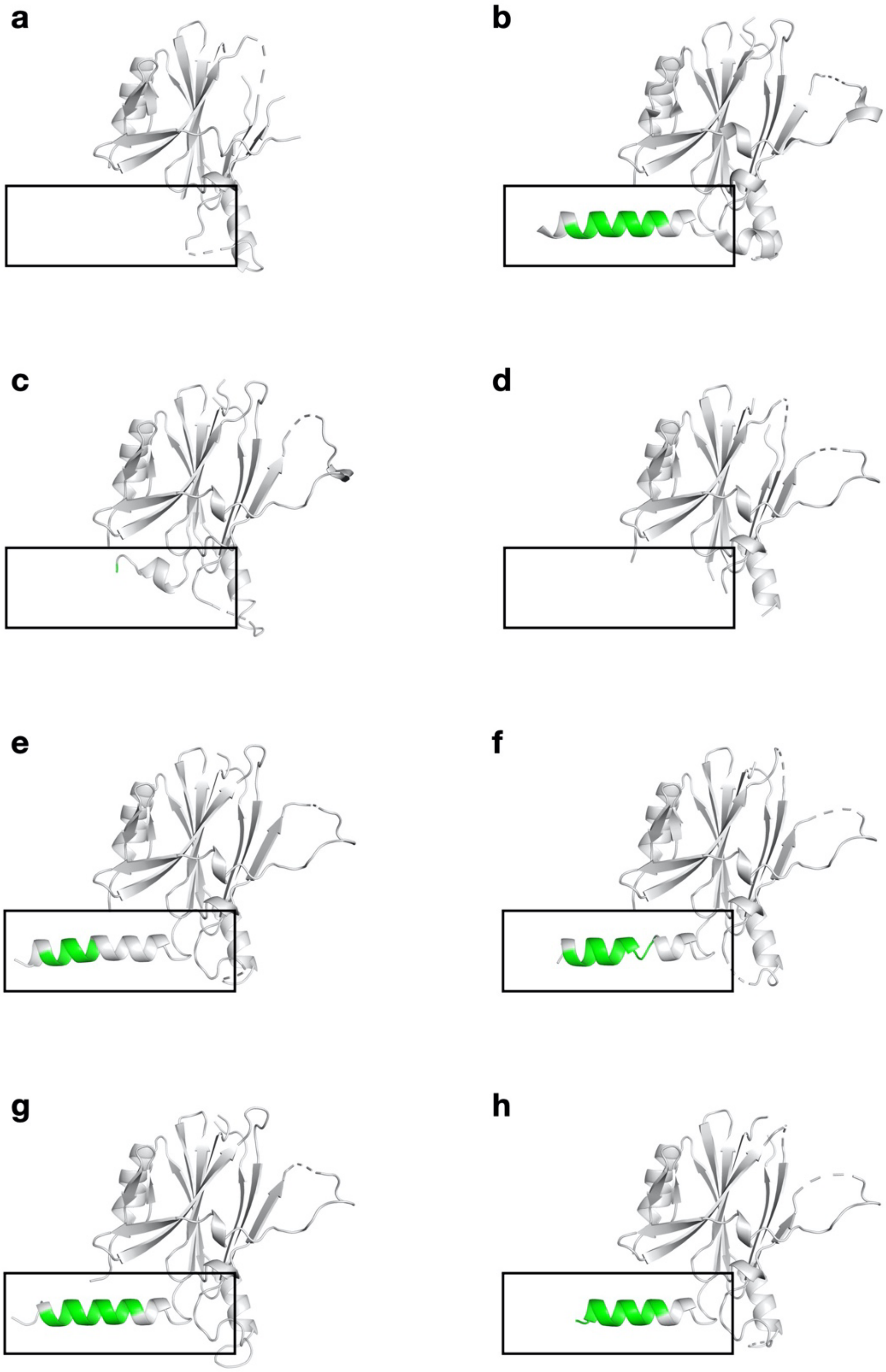


**Fig. S7** The crystal structures of (a) **c-Myc_1/PhC$_{\Delta14–24}$** (3.52 Å) (b) **c-Myc_1/PhC$_{\Delta15–25}$** (1.92 Å), (c) **c-Myc_2/PhC$_{\Delta15–25}$** (1.90 Å), and (d) **c-Myc_2/PhC$_{\Delta16–26}$** (2.29 Å), (e) **c-Myc_1/PhC$_{\Delta15–20}$** (2.55 Å), (f) **c-Myc_1/PhC$_{\Delta15–22}$** (2.55 Å), (g) **c-Myc_1/PhC$_{\Delta15–25}$_R151Q** (2.00 Å), (h) **c-Myc_1/PhC$_{\Delta15–25}$_R151K** (2.04 Å). Black squares show the position of the inserted c-Myc_1 and c-Myc_2 fragments. The green ribbon models in (b–h) show inserted c-Myc_1.

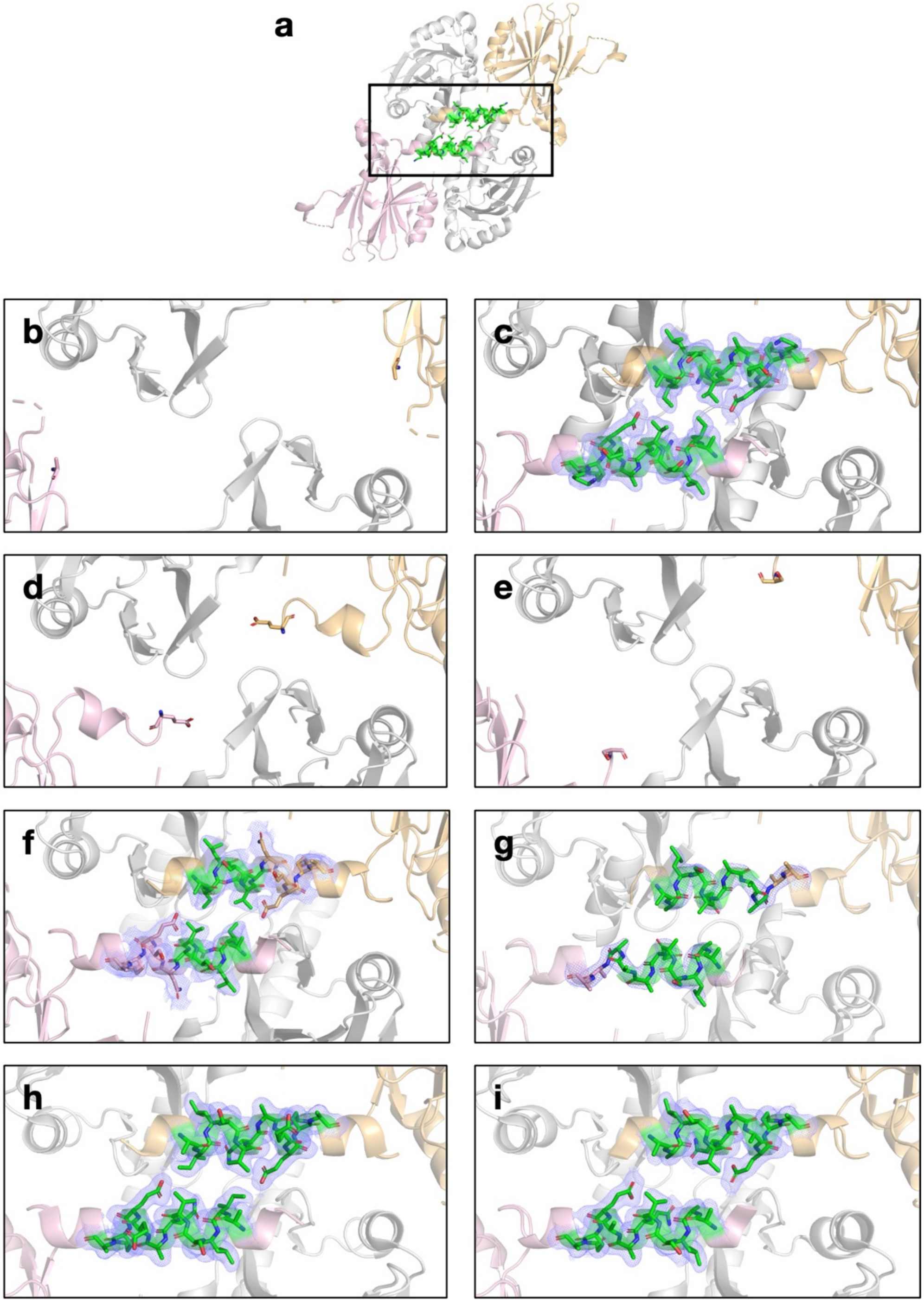


**Fig. S8** (a) The tetrameric structure of **c-Myc_1/PhC$_{\Delta15–25}$**. The close-up views of c-Myc fragments in (b) **c-Myc_1/PhC$_{\Delta14–24}$**, (c) **c-Myc_1/PhC$_{\Delta15–25}$**, (d) **c-Myc_2/PhC$_{\Delta15–25}$**, (e) **c-Myc_2/PhC$_{\Delta16–26}$**, (f) **c-Myc_1/PhC$_{\Delta15–20}$**, (g) **c-Myc_1/PhC$_{\Delta15–22}$**, (h) **c-Myc_1/PhC$_{\Delta15–25}$_R151Q**, and (i) **c-Myc_1/PhC$_{\Delta15–25}$_R151K**. The black square in (a) shows the inserted **c-Myc_1$^{\Delta15–25}$** position. The green ribbon models show the c-Myc_1 fragments. The selected 2|Fo|−|Fc| electron density maps at 1.0σ are shown in blue.

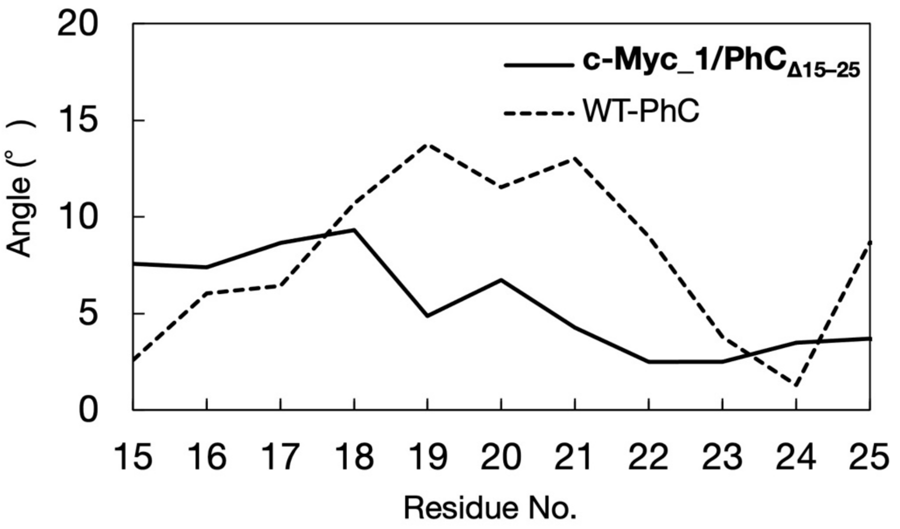


**Fig. S9** The local bending analysis of the H1 regions of **c-Myc_1/PhC$_{\Delta 15\text{-}25}$** and WT-PhC. The bending angle of a residue which residue No. is (n) is defined as the angle between two axes of the local helix of 4 residues at (n-3, n-2n n-1, n) and (n, n+1, n+2, n+3). The calculations were performed by the HELANAL algorithm.(10)

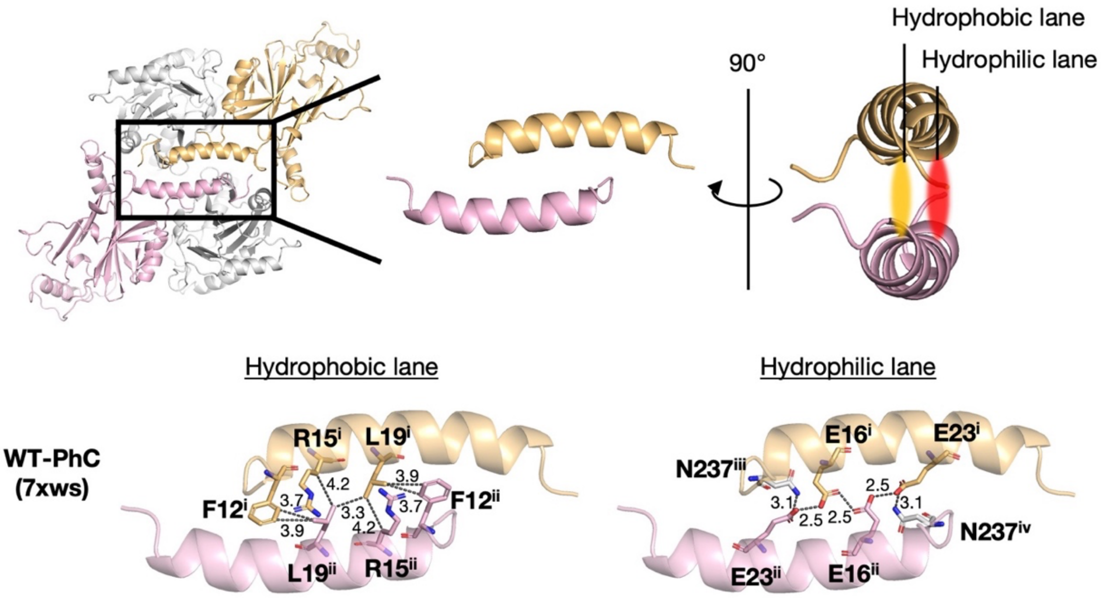


**Fig. S10** Inter-helix interaction of two-fold interface of H1s of WT-PhC. The cut-off distances of noncovalent interactions are 3.5 Å for hydrogen bonds, and 4.2 Å for hydrophobic interactions.(11, 12)

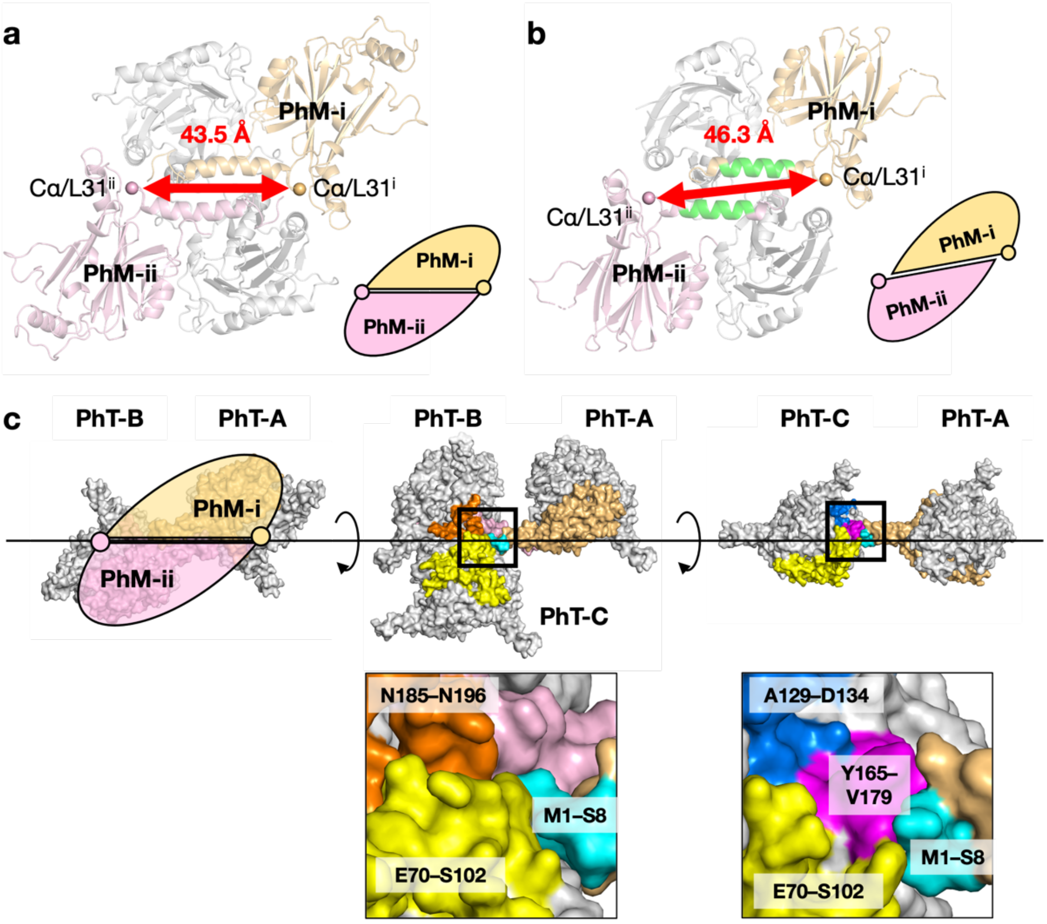


**Fig. S11** Comparison of the molecular packing in WT-PhC and **c-Myc_1/PhM**$_{\boldsymbol{\Delta 15\text{–}25}}$. Structures of the 4-mer of (a) WT-PhC and (b) **c-Myc_1/PhM**$_{\boldsymbol{\Delta 15\text{–}25}}$. (c) Structures of the interfaces among PhT-A, PhT-B, and PhT-C in WT-PhC. The Cα atoms are shown as spheres. PhTs are shown as the surface models.

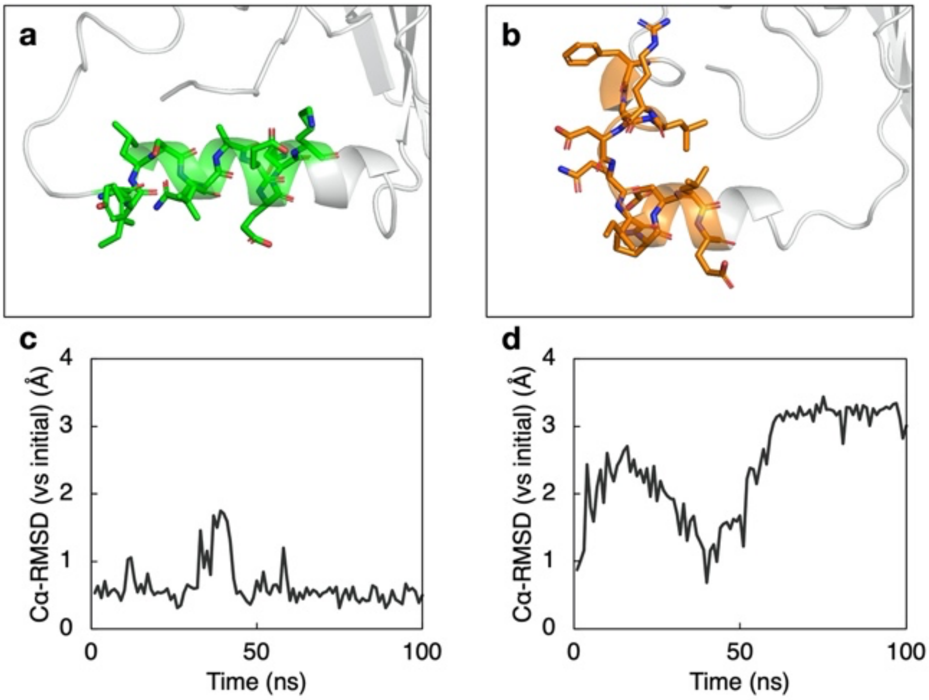


**Fig. S12** Analysis of conformations of c-Myc fragments in MD simulations of monomer. The structures at 100 ns in MD simulations of (a) **c-Myc_1/PhM$_{\Delta 15\text{–}25}$** and (b) **c-Myc_2/PhM$_{\Delta 15\text{–}25}$**. Time courses of the Cα-RMSD values of (c) c-Myc_1 and (d) c-Myc_2 from the initial structures. The ribbon models for c-Myc_1 in (a) and c-Myc_2 in (b) are colored in green and orange, respectively.

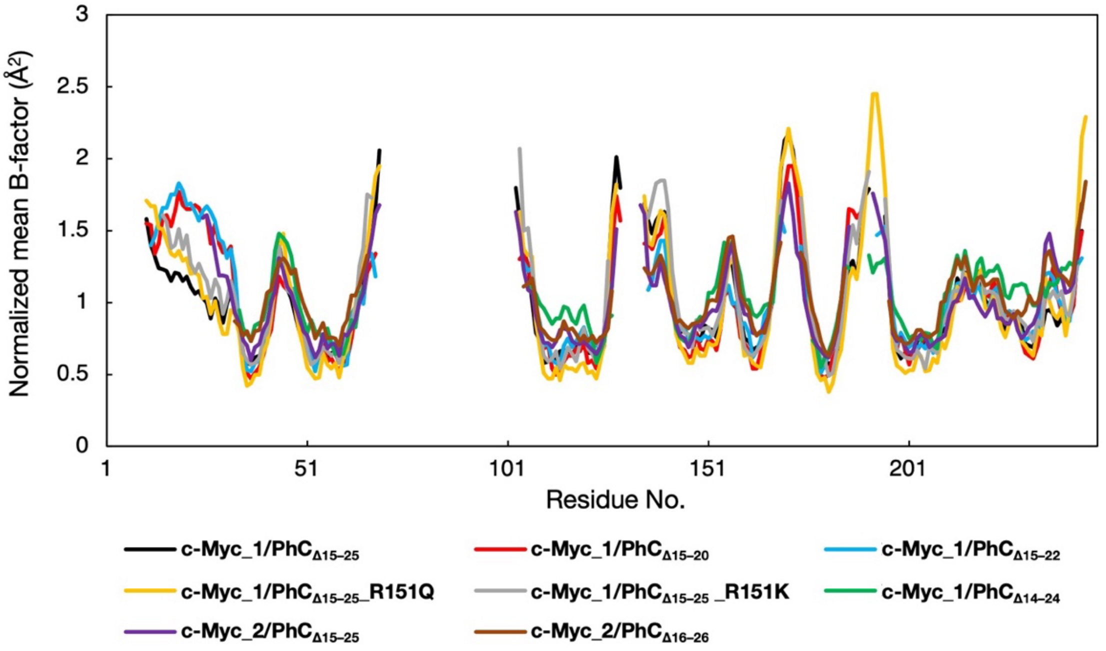


**Fig. S13** The spectrum of normalized mean B-factors (B') of main chain of **c-Myc_1/PhM$_{\Delta15\text{–}25}$, c-Myc_1/PhM$_{\Delta15\text{–}20}$**, **c-Myc_1/PhM$_{\Delta15\text{–}22}$**, **c-Myc_1/PhM$_{\Delta15\text{–}25}$_R151Q, c-Myc_1/PhM$_{\Delta15\text{–}25}$_R151K, c-Myc_1/PhM$_{\Delta14\text{–}24}$**, **c-Myc_2/PhM$_{\Delta15\text{–}25}$**, and **c-Myc_2/PhM$_{\Delta16\text{–}26}$**.

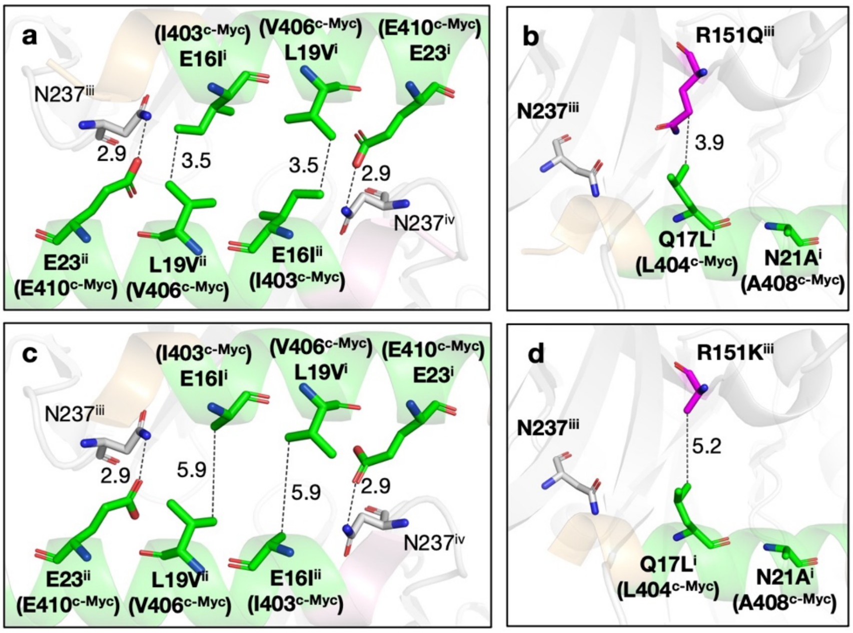


**Fig. S14** The intermolecular interactions networks of the two-fold interface of c-Myc_1 fragment of (a) **c-Myc_1/PhM$_{\Delta15\text{–}25}$_R151Q** and (c) **c-Myc_1/PhM$_{\Delta15\text{–}25}$_R151K**. Intermolecular interactions at R151$^{iii}$ of (b) **c-Myc_1/PhM$_{\Delta15\text{–}25}$_R151Q** and (d) **c-Myc_1/PhM$_{\Delta15\text{–}25}$_R151K**. All fragments of c-Myc_1 are colored green. R151Q$^{iii}$ in (b), and R151K$^{iii}$ in (d) are colored magenta. The cut-off distances of noncovalent interactions are 3.5 Å for hydrogen bonds, and 5.75 Å for hydrophobic interactions.(11, 12)

**Table S1** Rosetta energy analysis using Foldit.

| Mutant PhMs (H1 region) | Rosetta energy (E) of tetramer (kcal/mol) |
|---|---|
| WT-PhM | 903.5 |
| **c-Myc_1/PhM$_{\Delta 14–24}$** | 1522.1 |
| **c-Myc_1/PhM$_{\Delta 15–25}$** | 931.4 |
| **c-Myc_1/PhM$_{\Delta 16–26}$** | 1033.0 |

| Mutant PhMs (L1 region) | Rosetta energy (E) of trimer (kcal/mol) |
|---|---|
| WT-PhM | 650.6 |
| **c-Myc_1/PhM$_{\Delta 71–81}$** | 585.4 |
| **c-Myc_1/PhM$_{\Delta 72–82}$** | 605.4 |
| **c-Myc_1/PhM$_{\Delta 73–83}$** | 627.6 |

**Table S2** Crystallographic data of c-Myc_1 and c-Myc_2 fused PhCs.

| | c-Myc_1/PhC$_{\Delta 14–24}$ | c-Myc_1/PhC$_{\Delta 15–25}$ | c-Myc_2/PhC$_{\Delta 15–25}$ | c-Myc_2/PhC$_{\Delta 16–26}$ | c-Myc_1/PhC$_{\Delta 15–20}$ | c-Myc_1/PhC$_{\Delta 15–22}$ | c-Myc_1/PhC$_{\Delta 15–25}$_R151Q | c-Myc_1/PhC$_{\Delta 15–25}$_R151K |
|---|---|---|---|---|---|---|---|---|
| **Data collection** | | | | | | | | |
| Space group | *I*23 | *I*23 | *I*23 | *I*23 | *I*23 | *I*23 | *I*23 | *I*23 |
| Cell dimensions *a*=*b*=*c* (Å) | 107.91 | 105.97 | 106.40 | 106.75 | 106.10 | 105.14 | 106.59 | 106.53 |
| Resolution range (Å) | 50–3.52 (3.73–3.52) | 50–1.92 (1.99–1.92) | 50–1.90 (1.97–1.90) | 50–2.29 (2.43–2.29) | 50–2.55 (2.70–2.55) | 50–2.55 (2.70−2.55) | 50–2.00 (2.07–2.00) | 50–2.04 (2.11–2.04) |
| Observed reflections | 46,528 (7,153) | 1,335,868 (131,410) | 929,676 (93,397) | 994,943 (156,438) | 165,166 (26,276) | 252,308 (40,081) | 1,527,026 (148,820) | 753,098 (112,322) |
| Unique reflections | 2,709 (432) | 15,252 (1,486) | 15,963 (1,583) | 9,266 (1,491) | 6,629 (1,024) | 6,468 (1,003) | 13,787 (1,365) | 12,981 (2,027) |
| Redundancy | 17.2 (16.6) | 87.6 (88.4) | 58.2 (59.0) | 107.4 (104.9) | 24.9 (25.7) | 39.0 (40.0) | 2.0 (2.0) | 58.0 (55.4) |
| CC(1/2) | 98.8 (61.1) | 99.5 (73.9) | 100.0 (74.0) | 99.9 (63.8) | 98.2 (59.7) | 99.6 (64.3) | 99.9 (74.6) | 99.9 (65.8) |
| I/σ(I) | 13.8 (1.8) | 12.0 (1.3) | 24.3 (1.5) | 26.54 (1.17) | 7.35 (1.38) | 12.7 (1.2) | 26.1 (1.7) | 19.4 (1.3) |
| Completeness (%) | 99.5 (100.0) | 99.7 (98.2) | 99.9 (100.0) | 100.0 (100.0) | 100.0 (100.0) | 100.0 (100.0) | 99.9 (99.9) | 99.9 (100.0) |
| **Refinement** | | | | | | | | |
| Resolution (Å) | 34.12–3.52 | 37.47–1.92 | 37.62–1.90 | 33.76–2.29 | 43.32–2.55 | 33.25–2.55 | 37.69–2.00 | 37.66–2.04 |
| Number of reflections | 2,698 | 15,161 | 15,958 | 9,264 | 6,629 | 6,464 | 13,784 | 12,979 |
| *R*-factor (%) | 27.15 | 20.18 | 20.53 | 24.31 | 20.05 | 21.59 | 19.94 | 22.19 |
| Free *R*-factor (%) | 34.47 | 24.53 | 23.63 | 30.96 | 28.01 | 26.85 | 24.97 | 26.48 |
| R. m. s. deviations | | | | | | | | |
| Bond lengths (Å) | 0.003 | 0.008 | 0.009 | 0.009 | 0.008 | 0.008 | 0.008 | 0.008 |
| Bond angles (°) | 0.63 | 1.00 | 1.08 | 1.04 | 0.99 | 0.99 | 0.82 | 0.93 |
| Ramachandran plot (%) | | | | | | | | |
| favoured | 91.30 | 98.36 | 97.63 | 94.04 | 97.24 | 95.18 | 96.74 | 97.63 |
| allowed | 7.97 | 1.64 | 1.78 | 4.64 | 2.76 | 4.82 | 3.26 | 2.37 |
| outlier | 0.72 | 0.00 | 0.59 | 1.32 | 0.00 | 0.00 | 0.00 | 0.00 |
| PDB ID | − | 8J2Q | − | − | 8WLF | 8WLG | 8X8V | 8X8S |

Values in parentheses are for the highest-resolution shell.
